# Protection promotes energetically efficient structures in marine communities

**DOI:** 10.1101/2022.06.02.494503

**Authors:** Andrea Tabi, Luis J. Gilarranz, Spencer A. Wood, Jennifer A. Dunne, Serguei Saavedra

**Affiliations:** Institute for Cross-Disciplinary Physics and Complex Systems (IFISC), Consejo Superior de Investigaciones Científicas (CSIC) and University of Balearic Islands, 07122 Palma de Mallorca, Spain; School of Biological Sciences, University of Canterbury, Private Bag 4800, Christchurch 8140, New Zealand; Te Pūnaha Matatini, Centre of Research Excellence in Complex Systems, New Zealand; Department of Aquatic Ecology, Eawag (Swiss Federal Institute of Aquatic Science and Technology), Überlandstrasse 133, 8600, Dübendorf, ZH, Switzerland; College of the Environment, University of Washington, Seattle, WA 98195, USA; Santa Fe Institute, Santa Fe, NM 87501, USA; Department of Civil and Environmental Engineering, MIT, 77 Massachusetts Av., Cambridge, MA 02139, USA

**Keywords:** metabolic scaling, causal inference, community structure, transfer efficiency, pertur-bations, protected areas

## Abstract

The sustainability of marine communities is critical for supporting many biophysical processes that provide ecosystem services that promote human well-being. It is expected that anthropogenic disturbances such as climate change and human activities will tend to create less energetically-efficient ecosystems that support less biomass per unit energy flow. It is debated, however, whether this expected development should translate into bottom-heavy (with small basal species being the most abundant) or top-heavy communities (where more biomass is supported at higher trophic levels with species having larger body sizes). Here, we combine ecological theory and empirical data to demonstrate that full marine protection promotes shifts towards top-heavy energetically-efficient structures in marine communities. First, we use metabolic scaling theory to show that protected communities are expected to display stronger top-heavy structures than disturbed communities. Similarly, we show theoretically that communities with high energy transfer efficiency display stronger top-heavy structures than communities with low transfer efficiency. Next, we use empirical structures observed within fully protected marine areas compared to disturbed areas that vary in stress from thermal events and adjacent human activity. Using a nonparametric causal-inference analysis, we find a strong, positive, causal effect between full marine protection and stronger top-heavy structures. Our work corroborates ecological theory on community development and provides a quantitative framework to study the potential restorative effects of different candidate strategies on protected areas.

**Preprint:** The manuscript [1] is deposited on bioRxiv (https://doi.org/10.1101/2022.06.02.494503).

## Introduction

Human activities and environmental change are accelerating rates of biodiversity loss from ecosystems worldwide [2, 3, 4]. Through impacts on the geographical distributions, population abundances, and body size of organisms, anthropogenic stressors such as climate change and harvesting can fundamentally alter community composition [3, 5, 6]. For example, functional coral reefs and marine ecosystems are critical for maintaining the biophysical processes that support fisheries and other ecosystem services that contribute to human well-being [7, 8, 9]. Yet, the loss of coral can occur because of thermal stress as well as land- and ocean-based human activities [10], which in turn can lead to cascading effects on entire reef-associated communities [11]. Because restoration and conservation efforts require interventions, it then becomes necessary to increase our understanding of cause-effect relationships (not just correlations) between disturbance and community composition.

It is hypothesized that less-disturbed communities will tend to develop more energetically-efficient systems (i.e., support more biomass per unit energy flow) [12, 13]. This hypothesis is based on both the Energetic Equivalence Hypothesis (EEH) [14], which states that the total energy used by different species tends to be constant (i.e., independent of body size), and Metabolic Scaling Theory (MST) [15], which is built on the sub-linear scaling of standard metabolic rate with body size [16, 17]. Following MST, the power-law scaling exponent between body mass and biomass is therefore expected to be around 1*/*4 or less accounting for the inefficiency of transfer of energy across trophic levels [18]. In turn, population biomasses within ecosystems vary as a function of the species body size—commonly referred as differences in *community structure* [19, 20, 21]. Body size or mass is considered a “master trait” that scales with organisms’ physiology, regulating metabolic requirements [16], constraining feeding range [22], and shaping the trophic position of species in marine food webs due to energy transfers [23]. It is debated, however, whether less disturbed systems should translate into bottom-heavy structures (small basal species are the most abundant and large apex predators the least abundant) or top-heavy structures (more biomass can be supported at higher trophic levels with species having larger body size) [20, 24, 25, 26, 27]. Debates continue about these hypotheses because of the lack of feasible interventions that can be done to test theoretical predictions in marine communities. For example, deviations of community structures in marine communities from theoretical expectations have been explained by processes including [25, 26] complex predatory behavior (e.g., large predators feed on lower trophic levels or have wider diet width [20]), foraging of mobile consumers for energy subsidies provided by fish spawning grounds [28, 24, 29], increased rates of trophic energy flux due to warming [15], decreased body size due to higher temperatures [30], and noise in local sampling [31]. Yet, understanding the link between disturbance, efficiency, and structure is essential for determining the factors regulating the dynamics and sustainability of marine communities.

To address the debate between bottom- and top-heavy ecosystems, we need well-defined experiments that eliminate all sources of bias using randomized controlled trials and test the effectiveness of a given intervention [32]. Indeed, while observational data are designed to predict likely mechanisms or processes, they cannot establish cause-effect relationships, only associations [32, 33]. That is, following Reichenbach’s principle [34], if two variables are statistically related, then there exists a third variable that causally influences both (known as a confounding effect). Under specific contexts, this third variable can be one of the two variables, establishing a causal relationship between them. In this line, causal inference tools, such as path analysis or structural equation modeling [33], have been developed to obtain information about causes from observations. While extremely useful, these tools assume linearity or monotonicity in all the relationships, but many times this can be difficult to prove [32, 35]. Nevertheless, new advancements in nonparametric, causal, inference analysis allow us to investigate the structure and extent to which a likely cause can affect the probability that a given effect happens without many assumptions [32, 36].

As it is unfeasible to perform large-scale and controlled experiments of disturbance in marine communities, marine protected areas (MPAs) present unique observational opportunities to infer the causal relationship between protection from human disturbance and community structure in conjunction with differing levels of thermal stress and human activity. First, we use metabolic scaling theory [15] to establish theoretical predictions about the cause-effect relationship between protection from disturbance and structure of marine communities. Next, we use the empirical community structures observed within fully protected marine areas compared to disturbed areas across 299 geographical sites worldwide, comprising population data from 1,479 non-benthic marine species. Because no two communities are subject to the same internal [20, 29] (e.g., interspecific effects) and external conditions [37] (e.g., thermal stress), we follow a nonparametric causal-inference analysis [32, 36] to discover causal hypotheses and test the existence of a *genuine* causal relationship between marine protection and top-heavy structures. Finally, we discuss the implications of our results for the protection of marine communities and future avenues of research.

## Results

### Theoretical Analysis

To establish our theoretical predictions, we start by studying how changes in transfer efficiency (TE) across trophic levels affect the structure of marine communities. Specifically, we conduct a synthetic analysis based on metabolic scaling theory [15]. Following Ref. [19], we assume that size-based predation is responsible for the pathways of energy transfer in food webs from basal to higher trophic levels (see Methods for details). First, we randomly generate food web matrices based on the general niche model [38]. Second, using scaling relationships [39], for each community, we calculate average body masses for each species *i* as 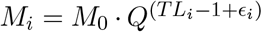, where *M*_0_ is a mass constant (here *M*_0_ = 1), *TL*_*i*_ corresponds to average trophic level, *Q* is the body-mass ratio across trophic levels (set to 10^3^), and random noise *ϵ*_*i*_. Thus, body mass increases with trophic level based on empirical observations [40, 24], so that the average predator-prey mass ratio ranges between 10 − 10^4^ in each community (Fig. 1A & C). Third, following Refs. [40, 20], we determine the transfer efficiency of each species *i* proportional to its body mass as 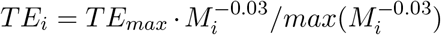 where the maximum trophic transfer efficiency (*TE*_*max*_) denotes the largest value in the community for basal species, so that transfer efficiency decreases with body mass (see supplementary Fig. S2 A). Lastly, following Ref. [15], we assume that the population biomass is proportional to average body mass in the form 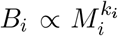, where *k*_*i*_ the individual biomass scaling coefficient (community structure). Following the energetic equivalence hypothesis with trophic transfer correction [18], the scaling coefficient is defined as 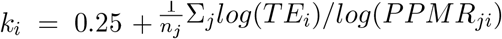 (see Methods for details). The predator-prey mass ratios (*PPMR*_*ji*_) were calculated for each species pair, which allows for incorporating trophic generality, especially for larger predators feeding on a wide variety of prey sizes [41, 42]. The community biomass scaling coefficient (*k*_*c*_) is calculated as the slope of the linear regression between log population biomasses (*B*_*i*_) and individual body sizes (*M*_*i*_) across all species *i*.

**Figure 1.**
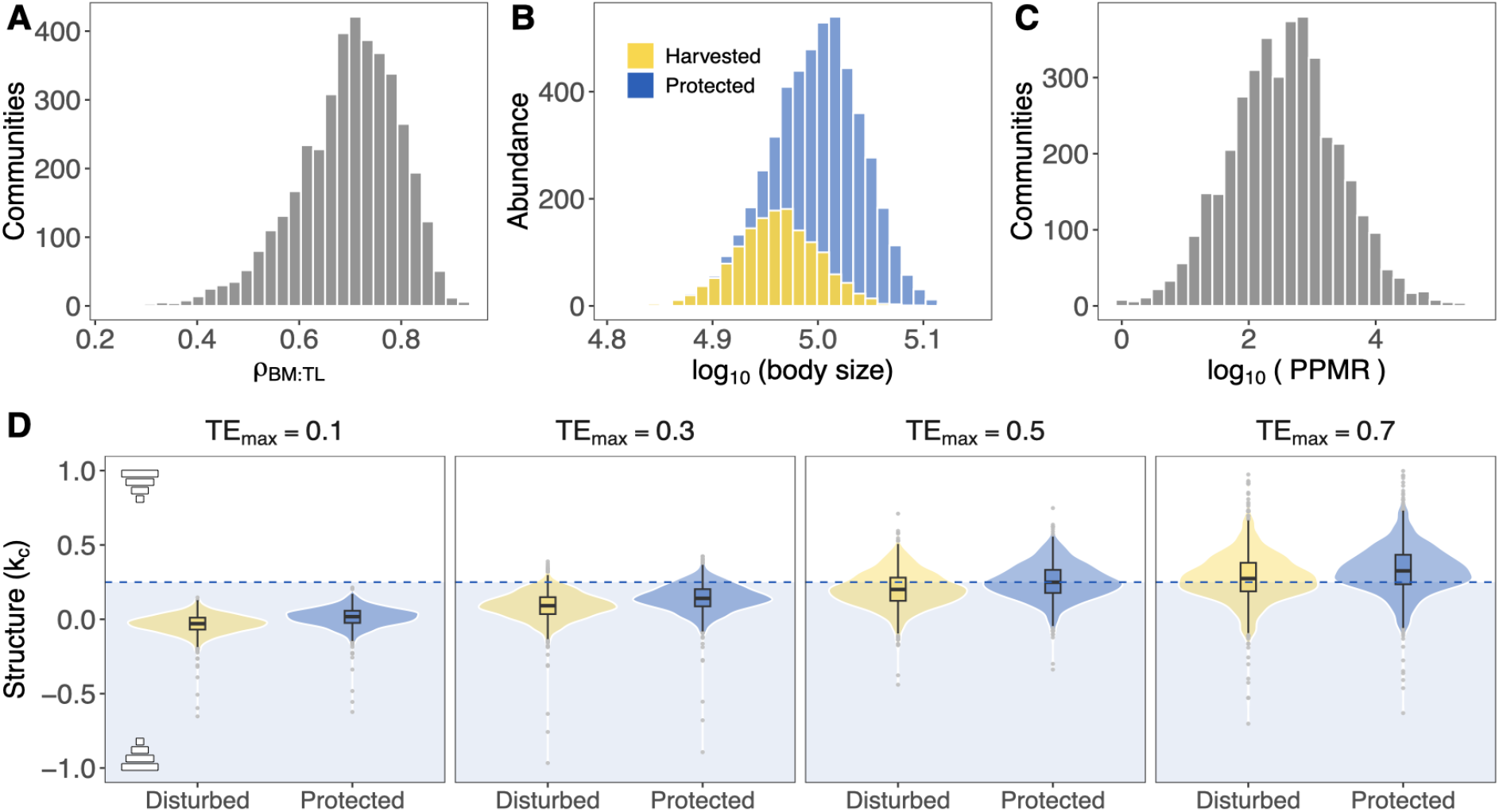
Theoretical predictions. Using Metabolic Scaling Theory (see Methods for details), Panel (A) depicts the distribution of Spearman’s rank correlation coefficients between average body size and average trophic level (*ρ*_*BS*:*T L*_). Panel (**B**) shows an example of how simulated selective harvesting affects the body size (measured as body mass) distribution of species. Specifically, selective harvesting is expected to reduce the number of individuals and average body size of larger-bodied species. Panel (C)shows the distribution of the median predator-prey mass ratios (PPMR) in each simulated community. Panel (**D**) shows that protected marine communities (blue boxplots) are expected to display stronger top-heavy structures than disturbed communities (yellow boxplots). Community structure is measured by the community scaling coefficient (*k*_*c*_), and higher values represent stronger top-heavy structures. Similarly, communities with higher transfer efficiency (*TE*_*max*_) display stronger top-heavy structures than communities with lower efficiency. Horizontal dashed line shows the expected scaling coefficient (*k*_*c*_ = 0.25) based on the Energetic Equivalence Hypothesis and lower values (light blue background) are the expected scaling coefficients accounting for trophic transfer efficiency.

To theoretically investigate the anthropogenic effect on community structure, we simulate a size-selective harvest of larger-bodied fish species as a potential source of disturbance [27] (see Methods for details). This selection effectively distorts body mass distributions by decreasing the biomass and the average body mass of the harvested species (Fig. 1B). After calculating the harvested biomasses, the disturbed community scaling coefficient is given as the slope of least square regression between log harvested biomasses and log harvested mean body sizes. Figure 1D shows that the median of *k*_*c*_ are higher for protected than disturbed communities due to the loss of biomass and decreased average body mass for higher trophic level species. However, differences are more pronounced when communities are characterized by higher transfer efficiencies. We showed that the theoretical expectations set by the EEH with trophic transfer correction (*k*_*c*_ ≤ 0.25) [18] can be largely exceeded by accounting for species having wide prey size spectrum including larger species (i.e., *PPMR <* 1) coupled with higher values of transfer efficiency. These theoretical results reveal that protected communities are expected to develop more energetically-efficient top-heavy structures, as developmental hypotheses suggest [13, 12].

### Empirical Analysis

To conduct our nonparametric causal inference analysis, we use observational data from marine reef-fish communities. These data comprise more than 1,500 fish species observations together with spatial, temporal, and climatic variables across 299 sampling sites worldwide from the Reef Life Survey database [43] (Fig. 2, Methods). For each sampling location, we compile data on whether the reef is within 10 km of a fully protected area (IUCN Category Ia: Strict Nature Reserve) as well as external conditions, including: whether it is associated with a coral reef within a 10-km radius, human population density (people per km^2^) within 25 km radius, as a proxy for human activity [44], and how frequently it experienced thermal stress anomalies (TSA) [45], as a measure of one climate-driven impact.

**Figure 2.**
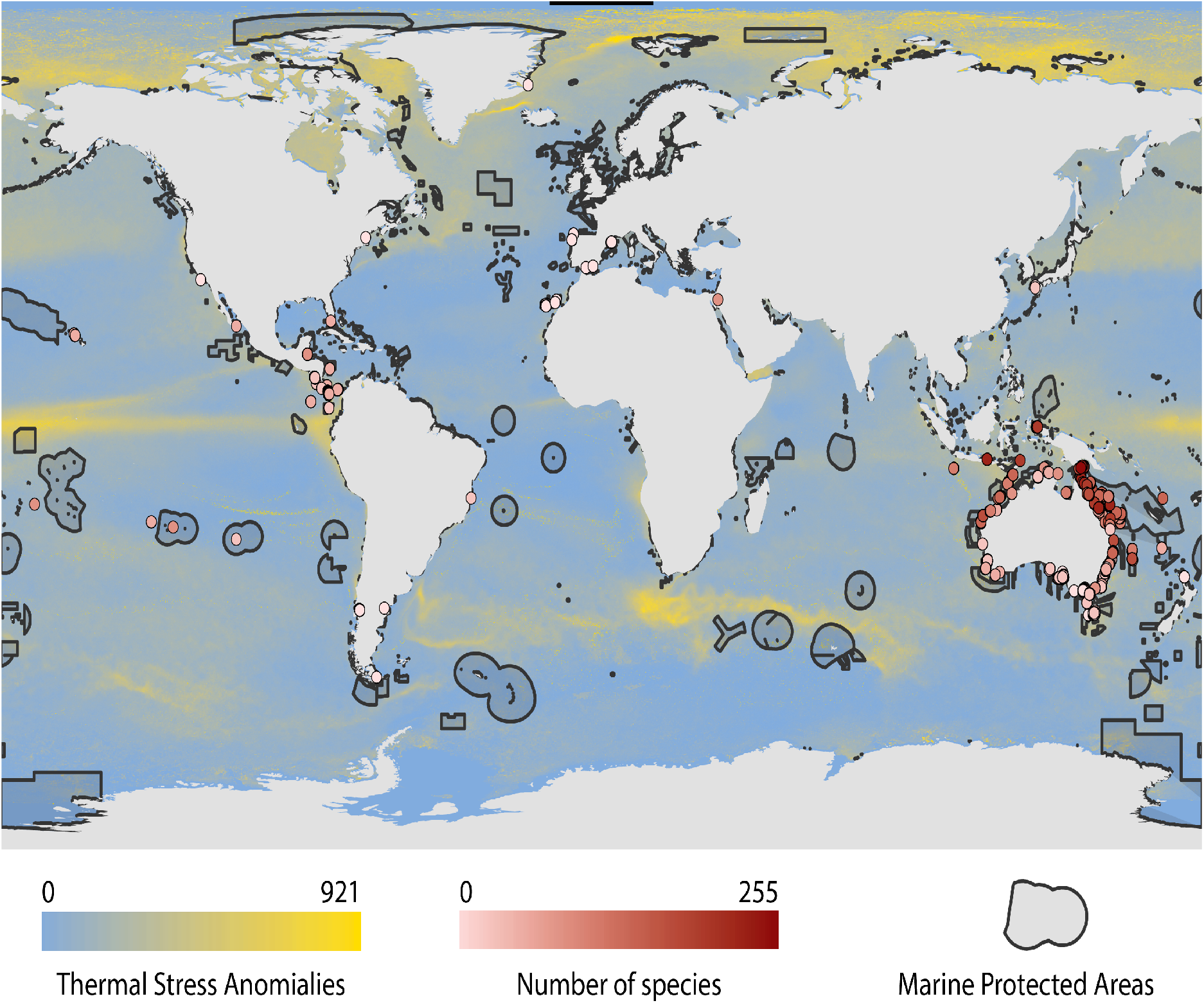
Global distribution of sampling sites and their attributes under our studied dataset. We consider only sampling sites (299 sites in total) in our analysis, which were surveyed more than once per year. Data compiled from the Reef Life Survey database [43] (see Methods for details). The color of the circles corresponds to the number of species observed at a given site. The background color corresponds to the thermal stress anomalies (TSA), which are calculated as the sum of all the values of TSA between 1982 and 2019, at which the average value of TSA was above 1 ^*°*^C. The black lines show the borders of reported Marine Protected Areas.

Community structure is traditionally measured by the power law exponent (*k*) between body sizes and biomass (or abundances) of species or trophic groups [20]. We calculate the empirical community scaling exponent 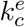as the slope of the least squares regression between log average body masses and log population biomasses of species for each community. The higher the values of 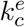, the stronger a community is characterized by a top-heavy structure. Because the theoretical biomass-size power law exponent is constrained to be *k* ≤ 0.25 unless predators have larger preys [24, 20]. However, the empirical biomass scale exponents largely exceeded the theoretical expectations (median value of *k*_*c*_ = 0.82) with higher average values in protected communities (Fig. 4). Empirical body mass distributions (measured as the community-weighted mean of body masses in each community) do not significantly differ in disturbed and protected communities (Fig. S3A), but the relationship between empirically measured average trophic level and average body mass is mostly positive in protected areas and it highly varies in disturbed areas (see supplementary Fig. S3B).

In order to investigate the existence of a causal relationship between marine protection and the structure of fish communities under the context of anthropogenic effects and climate change, we carried out a causal inference analysis [32]. First, we established a causal graph or hypothesis [32, 46]. Causal graphs can be constructed using expert knowledge or intuition. Alternatively, these graphs can be discovered using Inductive Causation (IC) [32]. We inferred the causal graph from empirical data using a standard IC algorithm (see Methods for details), which is based on conditional independence tests [32]. To both standardize and simplify our analysis, we transform all quantitative variables (community structure, MPAs, presence of coral reefs, thermal stress anomalies, and human density) into binary variables based on the median values. That is, values above the median are translated as *V* = 1, otherwise *V* = 0. Note that other variables are already binary by construction, such as the presence of protected areas and coral reefs. Figure 3 shows that the inferred causal graph supports the hypothesis that community structure and marine protection are causally related. Furthermore, the inferred graph supports the hypothesis that external drivers such as the presence of coral reefs, thermal stress anomalies, and human density influence the establishment of fully protected areas.

**Figure 3.**
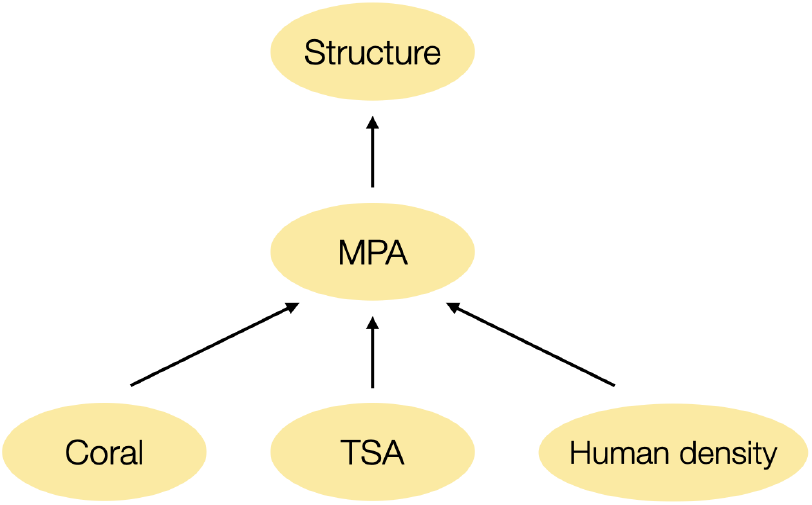
Inferred causal graph. Community structure is the direct cause of marine protection. Furthermore, TSA, coral reefs and human density affect the placement of MPAs. The causal graph was inferred using a causal discovery algorithm (Inductive Causation) based on conditional independence testing [32] (see Methods for more details).

**Figure 4.**
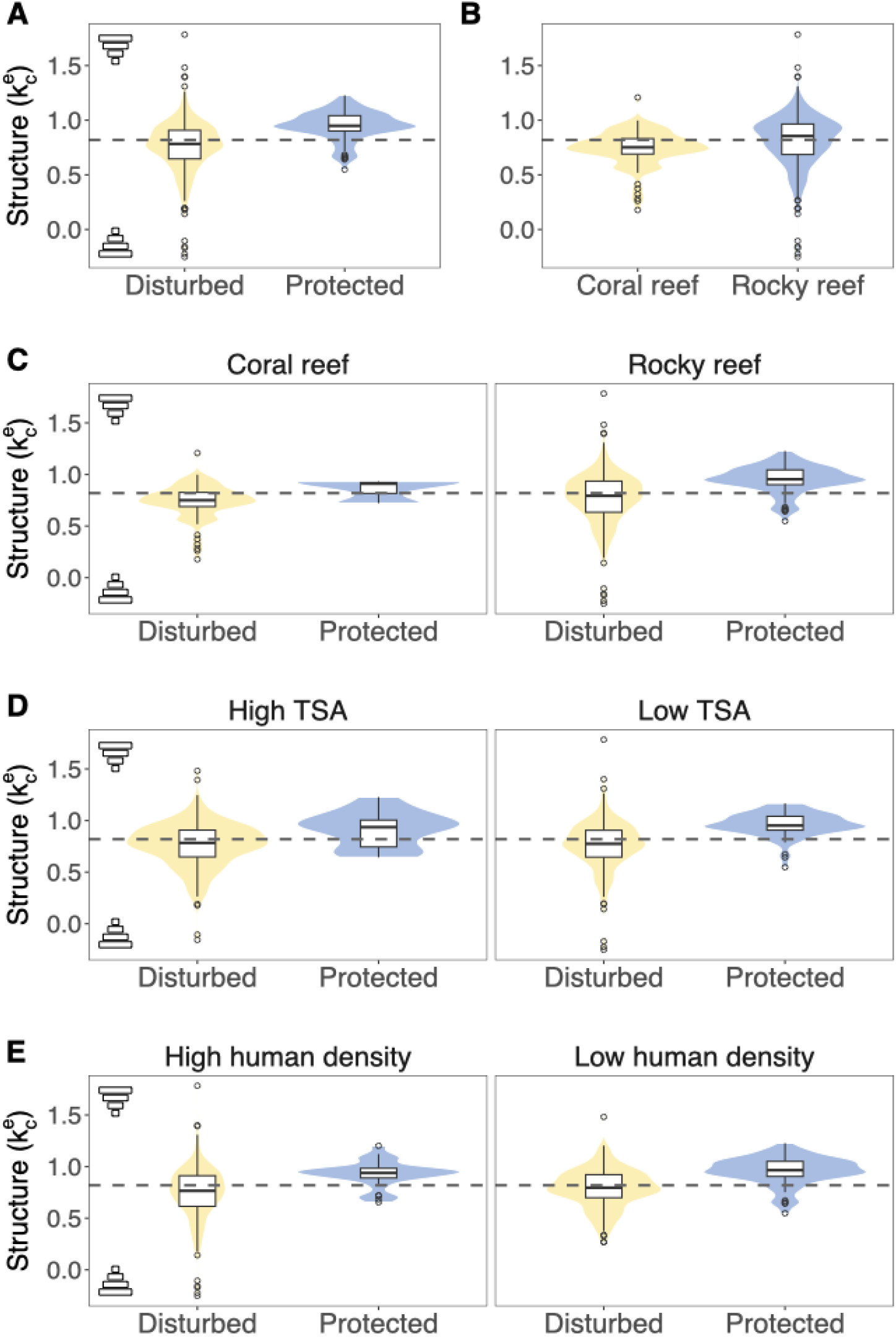
Distribution of community structures across different marine and geographical properties. The panels show the empirical distribution of community structures, measured as the regression coefficient 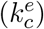 between log average individual body size and log population biomass. Higher values of 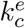 represent stronger top-heavy structures. Density: human density (people per km^2^) within 25 km radius following Ref. [44]. High and Low categories are Distributions separated by protected communities (MPAs under IUCN Category Ia) and disturbed communities. We transform all quantitative variables into binary variables based on the median values. That is, values above the median are translated as *V* = 1, otherwise *V* = 0. We refer to *V* = 1 (resp. *V* = 0) to *high* (resp. *low*) values. Note that some variables are already binary by definition, such as the presence or absence of protection and coral reefs.

Second, we tested whether the hypothesized relationship between marine protection and community structure is characterized by a *genuine* causal relationship and the extent of this relationship. In causal inference analysis [32], *genuine* causal relationships depict the strongest level of statistical support. This property requires the fulfilment of a statistical three-step criterion (see Methods for details). Because the nature of our causal hypothesis (Fig. 3), the existence of a *genuine* causal relationship between marine protection (*X*) and community structure (*Y*) can then be quantified as the average causal effect 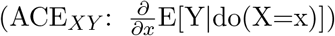 following the rules of *do*-calculus [32, 46] (see Methods for details). These rules allow us to translate (whenever possible) interventional conditional distributions *P* (*Y* = *y*|*do*(*X* = *x*)) into observational conditional distributions *P* (*Y* = *y*|*X* = *x*), such that ACE_*XY*_ = *P* (*Y* = 1|*do*(*X* = 1)) − *P* (*Y* = 1|*do*(*X* = 0))—put simply, the average causal effect is calculated as the probability of 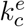 being higher than the median value in fully protected MPAs minus the probability of 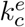being higher than the median value in disturbed areas.

Table 1 shows all six possible combinations under which it is possible to satisfy the three-step criterion necessary for inferring a *genuine* causal effect between protection from disturbance and community structure. Results show that all six combinations fulfilled the criterion for *genuine* causation between protection and community structure. Note that the greater the number of combinations, the stronger the statistical support for a *genuine* causal effect [32, 46]. Thus, following the relationships in Table 1 and the rules of *do*-calculus [32], the ACE between protection and structure 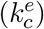 can be computed using the observational probabilities *ACE* = *P* (*k* = 1|Protection = 1) − *P* (*k* = 1|Protection = 0). Recall that we transform all variables into binary values (1: above median, 0: below median) and higher values of 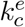 represent stronger top-heavy structures. Specifically, we find that fully protected areas directly increase by 43% (*ACE* = 0.431) the probability of observing fish communities with higher-than-average top-heavy structures. Indeed, Fig. 4 confirms that protected areas display stronger top-heavy structures than disturbed areas across any combination of the external variables (Table 1) required to fulfil the three-step criterion for a *genuine* causal relationship.

**Table 1:**
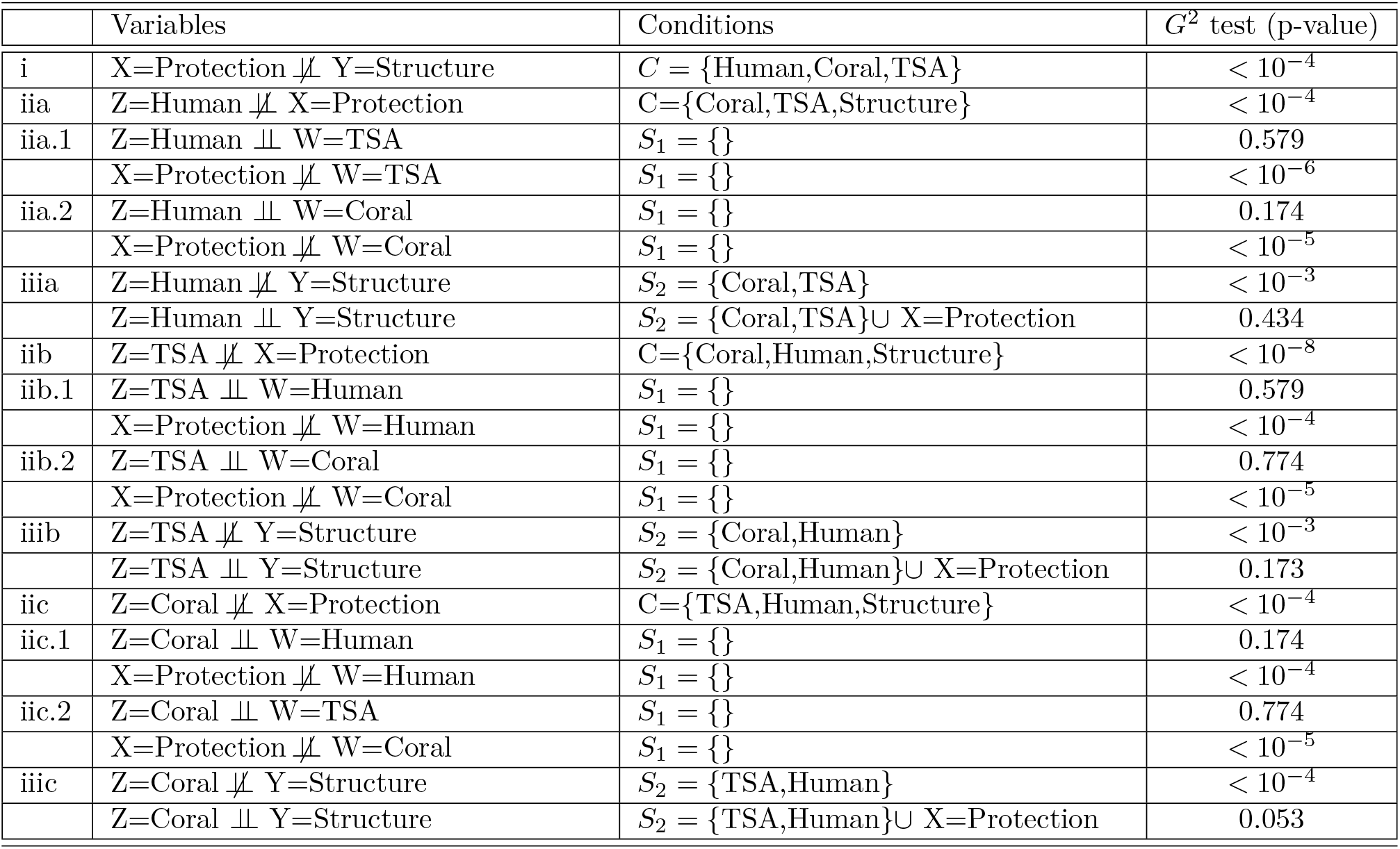
*Genuine* causal relationship between protection from harvesting and community structure. Here we test the criterion for inferring *genuine* causal relationship between protection and community structure under the context of TSA, human density and the presence of coral reefs. Following Ref. [32], we test the statistical 3-step criterion (i-iii) required to infer a *genuine* cause-effect relationship, the highest-level of causal inference that can be achieved (see Methods for details). Note that steps ii and iii have six alternative routes [32, 46]. That is, Route 1: i-iia-iia.1-iiia. Route 2: i-iia-iia.2-iiia. Route 3: i-iib-iib.1-iiib. Route 4: i-iib-iib.2-iiib. Route 5: i-iic-iic.1-iiic. Route 6: i-iic-iic.2-iiic. The larger the number of routes, the stronger the support. We use *G*^2^ test of independence [63]. We reject independency when the p-value < 0.05. ╨: independent, 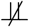: dependent

## Discussion

Our findings suggest that full marine protection (MPAs under IUCN Category Ia) regulates the structure of marine communities. Specifically, we have shown that protection in fully protected areas directly increases by 43% the probability that fish communities display a stronger top-heavy structure, relative to limits imposed by the environmental context, and thus supporting more biomass per unit of energy flow. Moreover, we show that top-heavy community structures in marine ecosystems are theoretically possible following the assumptions established by the Energetic Equivalence Hypothesis with trophic transfer correction [14, 15] incorporating trophic generality [41]. That is, the higher the transfer efficiency in marine communities, the stronger the magnitude and variation of top-heavy structures. By theoretically mimicking size-selective harvesting, we have also shown that harvested communities tend to develop more bottom-heavy structures compared to the unharvested state—consistent with empirical observations [27]. However, the body size distributions do not differ significantly between protected and disturbed communities, but the relationship between trophic level and average species body mass is largely positive in protected areas and highly variable in disturbed areas (Fig. S3) potentially due to overfishing [47]. In turn, other human activities, such as chemical pollution or habitat destruction, can reduce transfer efficiency through affecting nutrient availability [48]. Using fully protected areas, we have corroborated the existence of a positive *genuine* cause-effect relationship between full marine protection and top-heavy structures. The close match between our theoretical predictions and empirical findings supports the hypothesis that less disturbed ecosystems tend to be more energetically efficient [12, 13]. In regards to conservation and restoration efforts, our results open up the possibility to move from correlation to causation processes.

It is worth mentioning the scope and limitations of our work. Our theoretical model based on metabolic scaling relationships provides qualitative predictions regarding the shape of community structure, in terms of population biomass distribution across body masses and its association with trophic transfer efficiency. The shape of community structure carries information about ecological processes that potentially allow us to predict future community responses to disturbance and other types of environmental changes. While we were not able to calculate the trophic transfer efficiency in empirical communities directly, the similarity of our theoretical results to observed structural patterns strongly suggests that protected communities exhibit more energetically efficient structures compared to disturbed communities. Yet, the link between community structure and trophic transfer efficiency in empirical settings should be further investigated. Indeed, multiple processes can impact the efficiency of energy transfer in marine communities, such as different temperature sensitivity of metabolism across trophic levels, resource availability, and quality or non-predatory fluxes of organic material [49]. In fact, it is estimated that transfer efficiency varies widely between 1—52% across different regional and environmental contexts [49]. Our results point towards a large impact of transfer efficiency on community structure and composition, highlighting an important dynamic that has been understudied [50]. Additionally, in our theoretical analysis, we showed that predator-prey mass ratios together with transfer efficiency can generate a huge variety of community structures without violating the assumptions of EEH with trophic transfer correction. Most importantly, having a higher transfer efficiency alone can lead to top-heavy structures. For example, it has been shown that the presence of both large generalist predators and gigantic secondary consumers that feed much lower in the trophic web than predicted by size alone can lead to top-heavy structures [20]. However, other mechanisms such as spatial energy subsidies [24] can also contribute to reshaping community structure, therefore, more detailed information about predator-prey interactions is needed to separate the different mechanisms affecting community structure [51].

As human density and cumulative impacts in coastal areas increases [52, 53], and thermal stress anomalies become more frequent due to climate change [54, 5], it becomes increasingly important to sustain the function of marine communities [55]. While we have not studied the recovery of communities to a specific restoration baseline (which remains highly debated [56]), our results do point towards a strong, positive, *genuine* causal effect of full marine protection and the structure and efficiency of fish communities. Therefore, we believe that our theoretical and nonparametric methodologies can be used as a quantitative framework to study and guide experimental work focused on measuring the effect of potential interventions on relevant reference states of ecological communities in general.

## Methods

### Data

We analyzed 479 sampled communities from 299 sites (Fig. 2) from the Reef Life Survey database [43] comprising population data from more than 1,500 non-benthic marine species with individual body size information. Body size was measured as body mass and data were aggregated by year. We included only sampling sites in our analysis, which were surveyed more than once per year [57]. This decision was based on the rarefaction analysis and Kolmogorov–Smirnov tests to assess the impacts of annual sampling effort on species richness (see supplementary Section S1 for more details). We collected weekly sea surface temperature (SST) from NOAA’s (National Oceanic and Atmospheric Administration) remote sensing database. We used the sum of TSA in our analysis, calculated as the weekly sea surface temperature (SST) minus the maximum weekly climatological SST. TSA was measured as the number of events when the average difference between weekly SST and the maximum weekly climatological was above 1^*°*^C between 1982 and 2019 [45]. The distribution of warm-water coral reef was obtained from UNEP-WCMC World Fish Centre database [58]. The information on marine protected areas was obtained from UNEP-WCMC and IUCN Protected Planet database [59]. The information on human population density was obtained from Gridded Population of the World [60]. The human population density was quantified as *humans/Km*^2^ in a 25-km radius around the sampling site [52]. Lastly, we used the regression coefficient (*k*) between log biomass and log of average body sizes as a measure of community structure. The higher the values of *k*, the stronger a community is characterized by a top-heavy structure. Because the theoretical power-law exponent (*k*) is constrained to be *k* ≤ 0.25 unless predators are on average smaller than their prey [20]. We found qualitatively similar results if we use the Spearman’s rank correlation coefficient (*ρ*) between biomass and average body sizes as a measure of community structure. Values closer to *ρ* = 1 (resp. *ρ* = −1) specify communities closer to a perfect top-heavy (resp. bottom-heavy) structure.

### Theoretical analysis

To carry out our theoretical analysis, we randomly generated food web matrices of 35 species (in order to match the results with the empirical median richness level) based on the general niche model [38]. Following Ref. [61], we set the connectance of each food web given by the function of the number of species as *C* = *𝒮* ^−0.65^. Second, using scaling relationships [39], for each community, we calculated average individual body masses for each species *i* as 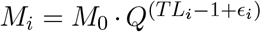, where *M*_0_ is a mass constant (here *M*_0_ is assumed to be 1), *TL*_*i*_ corresponds to average trophic level if each species, *Q* is the average body mass ratio across trophic levels set to 10^3^ (different values yield qualitatively similar results), and random noise *ϵ* ∼ *N* (0, 1). The predator-prey mass ratios (*PPMR*_*ji*_) were calculated for each species pairs based on the average body masses (*M*_*i*_*/M*_*j*_). The median predator-prey mass ratio ranges around 10 − 10^4^ in each community (Fig. 1A) corresponding to observations. Third, following Refs. [20, 40], we estimated the transfer efficiency of each species *i* proportional to its body mass as 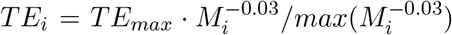, where *TE*_*max*_ is a scaling constant, which sets the maximum value of transfer efficiency of the community for the lowest trophic level. Fourth, to systematically investigate the effect of trophic transfer efficiency, we varied the scaling constant of transfer efficiency *TE*_*max*_ ∈ (0, 1)— higher values lead to higher efficiency. Fifth, following Ref. [15], we assumed that biomass is proportional to average body size in the form 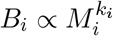, where the individual biomass scaling coefficient is defined as 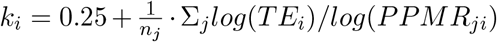. The community scaling coefficient (*k*_*c*_) was estimated as the slope of the least square regression between log biomass (*B*_*i*_) and log body masses (*M*_*i*_). Due to the properties of log ratios in the equation, we set *log*(*T E*_*i*_) if *PPMR*_*ij*_ *>* 1 and *log*(1 − *TE*_*i*_) if *PPMR*_*ij*_ *<* 1 in order to conserve the correct interpretation of increasing transfer efficiency. To theoretically investigate the potential effect of protection on community structure, we assumed a size-selective harvest of large fish species [27] as the source of disturbance. Following Ref. [62], size-selective harvest affects species in two ways; it decreases the average individual body mass and reduces the number of individuals. Thus, in each simulation, we set the fraction of species harvested to 40% (different percentages yield qualitatively similar results) and we determined the identity of harvested species from the community by randomly sampling where we assigned higher probability to larger fish species to be selected. As a next step, we randomly sampled the level of harvest for each fished species (*r*_*i*_) from a uniform distribution (*U* [0.3, 1]), where we set the minimum amount of removal at 30%. Finally, we calculated the harvested community biomass scaling coefficient 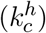 as the slope of the least square regression between log harvested population biomasses 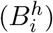 and log harvested individual body sizes 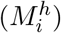.

### Causal discovery with Inductive Causation

The IC algorithm takes as input a stable probability distribution generated by some underlying causal structure and outputs a *pattern* (partially directed graph). The algorithm follows three major steps: 1. For each pair of variables 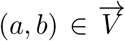, search for a set *S*_*ab*_ such that (*a* ╨ *b*|*S*_*ab*_). Construct an undirected graph *G* such that vertices (*a, b*) are connected with an edge if and only if no set *S*_*ab*_ can be found. 2. For each pair of non-adjacent variables (*a, b*) with a common neighbor *c*, check if *c* ∈ *S*_*ab*_. If it is then continue. If it is not, then orient *a* → *c* ← *b*. 3. In the partially directed graph that results, orient as many of the undirected edges as possible subject to two conditions (i) any alternative orientation would yield a new structure, or (ii) any alternative orientation would yield a directed cycle [32].

### *Genuine* causal relationship

Following Ref. [32], the subsequent statistical three-step criterion needs to be fulfilled in order to establish a *genuine* causal effect of random variable *X* on random variable *Y*. (i) *X* has to be statistically dependent on *Y* under a context *C* (set of additional variables). There must be a potential cause *Z* of *X*. This is true if *Z* and *X* are statistically dependent under context *C*, there is a variable *W* and context *S*_1_ ⊆ *C* such that *Z* and *W* are statistically independent, and *W* and *X* are statistically dependent. (iii) There must be a context *S*_2_ ⊆ *C* such that variables *Z* and *Y* are statistically dependent but statistically independent under the context *S*_2_ ∪ *X*. This 3-step criterion assumes that measured variables are affected by mutually independent, unknown, random variables.

### Rules of *do*-calculus

For readers’ convenience, here we write the three rules of *do*-calculus [32]. Let *G* be a DAG associated with a causal model and let *P* stand for the probability distribution induced by that model. Let 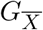 denote the graph obtained by deleting from *G* all arrows pointing to nodes in *X*. Likewise, 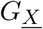denotes the graph obtained by deleting from *G* all arrows emerging from nodes in *X*. Finally, let *Z*(*W*) denote the set of *Z*-nodes that are not ancestors of any *W* -node. For any disjoint subset of variables *X, Y, Z* and *W*, we have the following three rules. Rule 1 (insertion/deletion of observations): 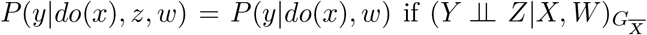. Rule 2 (action/observation exchange): 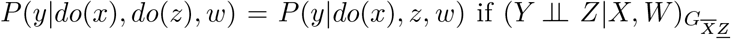. Rule 3 (insertion/deletion of actions): 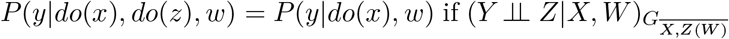. Note that ⊥: independent and 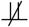⊥: dependent.

## Acknowledgments

A.T. was supported by the Spanish State Research Agency, through the Maria de Maeztu Program for Units of Excellence in R&D (MDM-2017-0711) and Te Pūnaha Matatini, a Centre of Research Excellence funded by the Tertiary Education Commission, New Zealand. L.J.G. was supported by the Swiss National Science Foundation Ambizione Fellowship, PZ00P3 185951. S.W. and J.D. were supported by the Santa Fe Institute. S.S. was supported by MIT Sea Grant College Program.

## Supplementary Information for

### S1 Sampling effort

We also showed that results can be dependent on the number of annual sampling events. The seasonal variations in fish populations affect population dynamics, which causes temporal changes in composition [64]. Therefore, we conducted a rarefaction analysis on the entire dataset to assess the effect of sampling effort on species richness at a given location and time (year). Figure S1 shows that reliable metrics about the communities can be produced only when the sampling effort is more than one time per year. Moreover, our rarefaction analysis concluded that increased sampling effort (*>* 1 per year) provides a more reliable description of the composition of communities (number of species: Figure S1, abundance of species: Table S1).

**Supplementary Figure S1:**
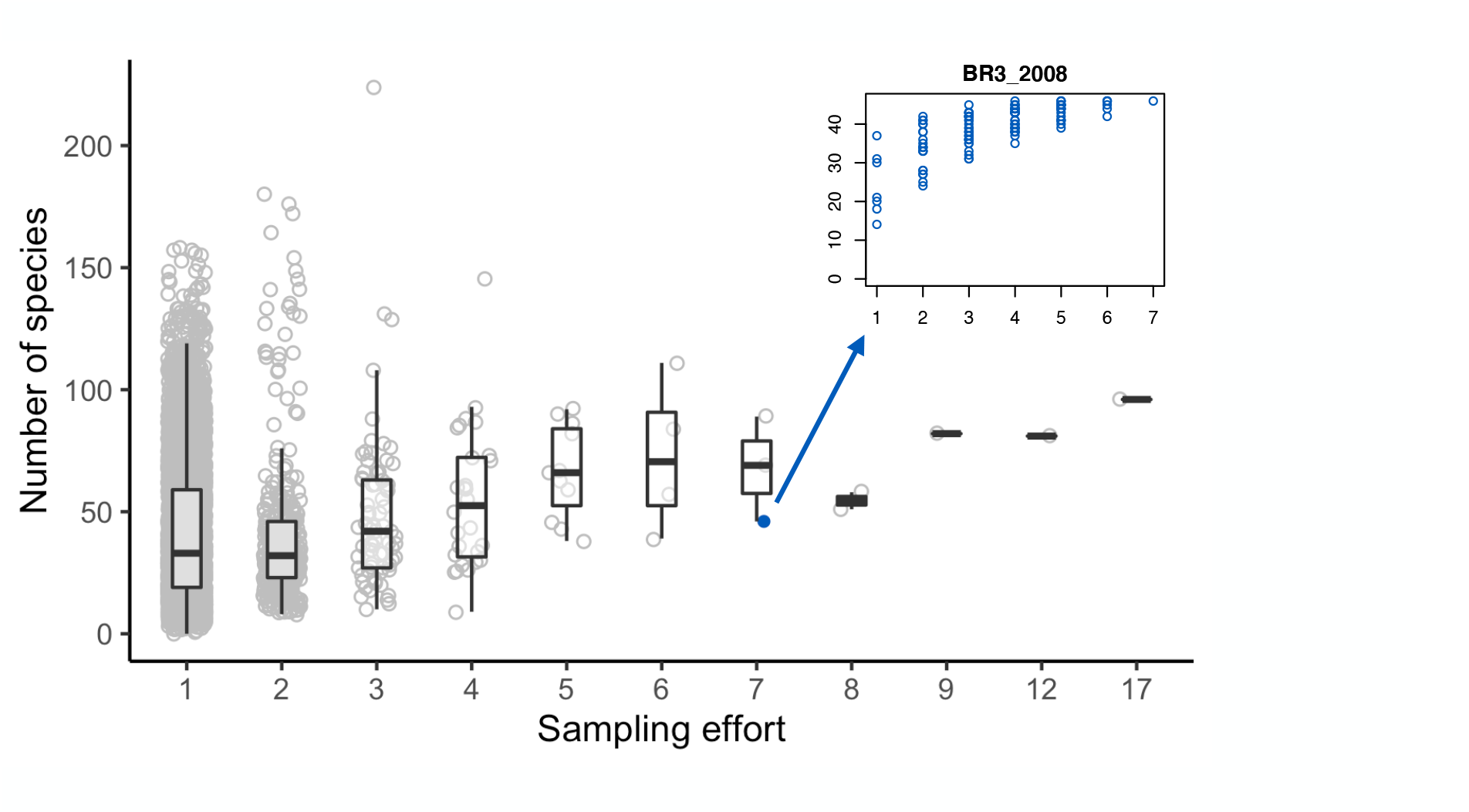
Sampling effort per year and an example of a rarefaction curve. The grey circles represent communities aggregated in a given location across a year. The majority of communities were sampled once per year, and only a small fraction of the aggregated communities were sampled more than once a year (sampling effort *>* 1). For communities sampled more than once in a given year, we conducted a rarefaction analysis to estimate the effect of sampling effort on species richness by resampling communities and then plotting the number of species in each constructed community against sampling effort (an example shown in the top right panel).

### S2 Supplementary Table

**Supplementary Table S1:**
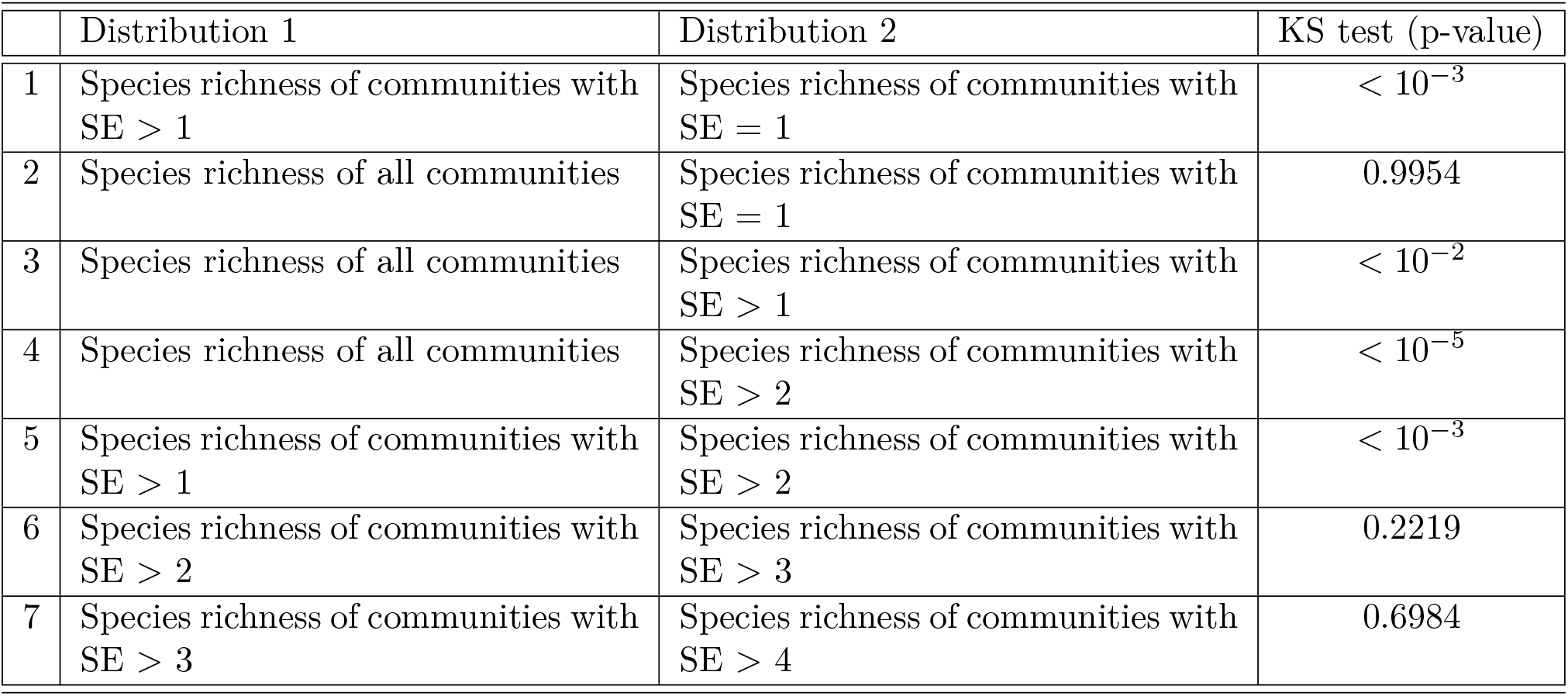
We used the Kolmogorov–Smirnov test to compare the empirical distributions of species richness across communities with different levels of sampling effort. Two samples are considered not drawn from the same distribution if the p-values *< α*, where we set *α* = 0.05.

|  | Distribution 1 | Distribution 2 | KS test (p-value) |
| --- | --- | --- | --- |
| 1 | Species richness of communities with SE > 1 | Species richness of communities with SE = 1 | $< 10^{-3}$ |
| 2 | Species richness of all communities | Species richness of communities with SE = 1 | 0.9954 |
| 3 | Species richness of all communities | Species richness of communities with SE > 1 | $< 10^{-2}$ |
| 4 | Species richness of all communities | Species richness of communities with SE > 2 | $< 10^{-5}$ |
| 5 | Species richness of communities with SE > 1 | Species richness of communities with SE > 2 | $< 10^{-3}$ |
| 6 | Species richness of communities with SE > 2 | Species richness of communities with SE > 3 | 0.2219 |
| 7 | Species richness of communities with SE > 3 | Species richness of communities with SE > 4 | 0.6984 |

### S3 Supplementary Figures

**Supplementary Figure S2:**
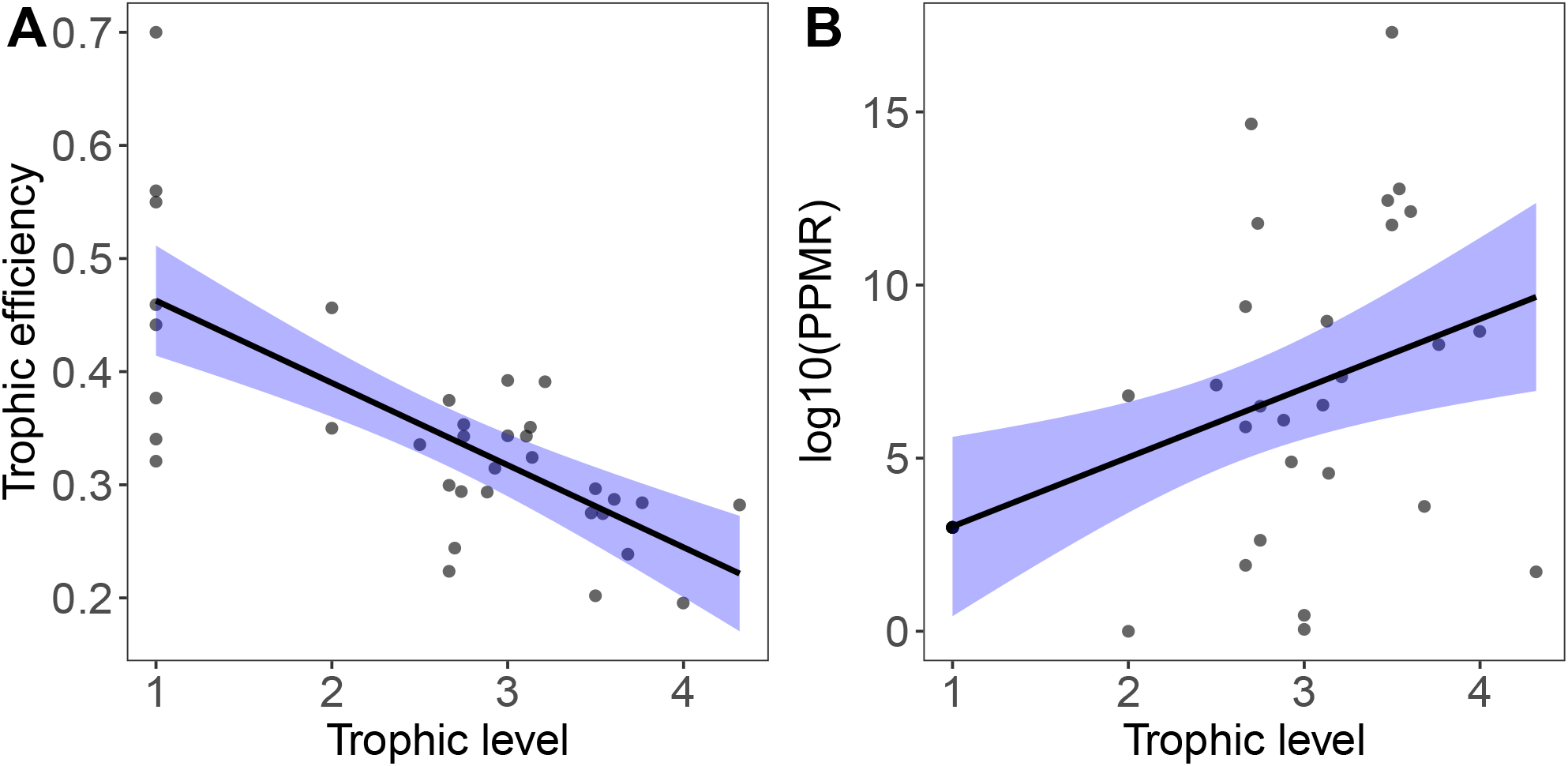
Theoretical distributions. Panel **A** shows an example of trophic efficiencies in a simulated community (*TEmax* = 0.7). On average trophic efficiency decrease with trophic level with some variation. Panel **B** shows that average predator-prey mass ratios (PPMR) increase with trophic level.

**Supplementary Figure S3:**
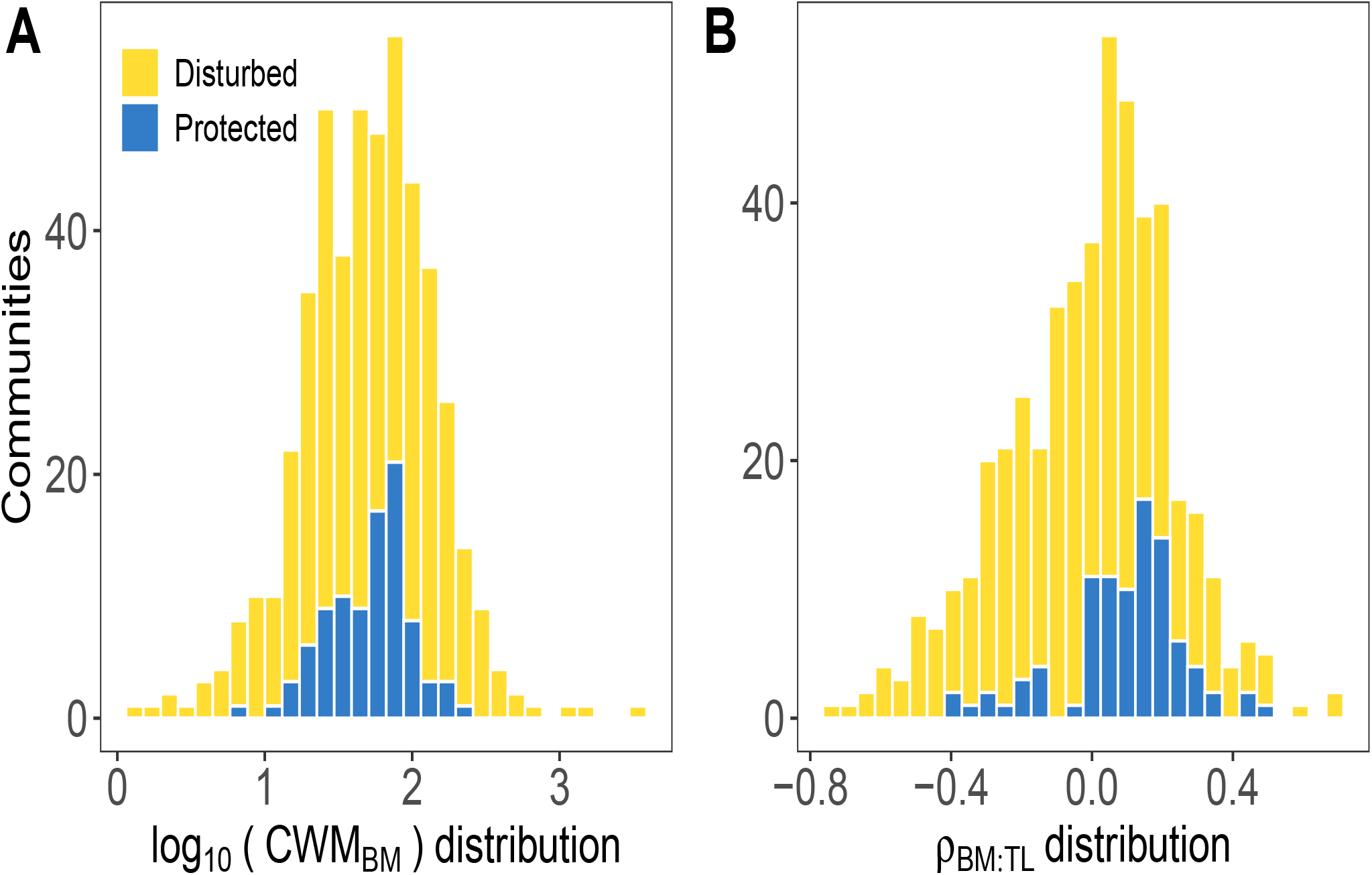
Empirical distributions. **A** Species body sizes were weighted by species abundance for each community. The average community-weighted mean of body masses (*CWM*_*BM*_) in protected and disturbed areas show similar values. **B** The distribution of Spearman’s rank correlation coefficient between average species body mass and trophic levels for empirical communities shows that empirical body masses do not always increase with trophic level. The average species trophic level was obtained from Fishbase database [65].

## References

[1] Tabi A, Gilarranz LJ, Wood SA, Dunne JA, Saavedra S (2023) Protection from harvesting promotes energetically efficient structures in marine communities.

[2] IPCC (2019) Special Report on the Ocean and Cryosphere in a Changing Climate [H.-O. Pörtner et al)]., Technical report.

[3] Halpern BS, et al. (2008) A Global Map of Human Impact on Marine Ecosystems. Science 319:948–952.

[4] Sala E, Knowlton N (2006) Global Marine Biodiversity Trends. Annual Review of Environment and Resources 31:93–122.

[5] Burrows MT, et al. (2011) The Pace of Shifting Climate in Marine and Terrestrial Ecosystems. Science 334:652–655.

[6] Poloczanska ES, et al. (2013) Global imprint of climate change on marine life. Nature Climate Change 3:919–925.

[7] Worm B, et al. (2006) Impacts of Biodiversity Loss on Ocean Ecosystem Services. Science 314:787–790.

[8] Palumbi SR, et al. (2009) Managing for ocean biodiversity to sustain marine ecosystem services. Frontiers in Ecology and the Environment 7:204–211.

[9] Spalding M, et al. (2017) Mapping the global value and distribution of coral reef tourism. Marine Policy 82:104–113.

[10] Mun∼iz-Castillo AI, et al. (2019) Three decades of heat stress exposure in Caribbean coral reefs: a new regional delineation to enhance conservation. Scientific Reports 9:11013.

[11] Ainsworth CH, Mumby PJ (2015) Coral–algal phase shifts alter fish communities and reduce fisheries production. Global Change Biology 21:165–172.

[12] Margalef R (1968) Perspectives in Ecological Theory (University of Chicago Press, Chicago).

[13] Odum EP (1969) The strategy of ecosystem development. Science 164:262–270.

[14] Nee S, Read AF, Greenwood JJD, Harvey PH (1991) The relationship between abundance and body size in British birds. Nature 351:312–313.

[15] Brown JH, Gillooly JF, Allen AP, M. V, West GB (2004) Toward a metabolic theory of ecology. Ecology 85:1771–1789.

[16] Kleiber M (1932) Body size and metabolism. Hilgardia 6:315–353.

[17] Elton CS (1927) Animal Ecology (University of Chicago Press, Chicago, IL).

[18] Brown JH, Gillooly JF (2003) Ecological food webs: High-quality data facilitate theoretical unification. Proceedings of the National Academy of Sciences 100:1467–1468.

[19] Sheldon RW, Prakash A, Sutcliffe Jr. WH (1972) The Size Distribution of Particles in the Ocean. Limnology and Oceanography 17:327–340.

[20] Woodson CB, Schramski JR, Joye SB (2018) A unifying theory for top-heavy ecosystem structure in the ocean. Nature Communications 9:1–8.

[21] Hatton IA, Heneghan RF, Bar-On YM, Galbraith ED (2021) The global ocean size spectrum from bacteria to whales. Science Advances 7:eabh3732.

[22] Andersen KH, et al. (2016) Characteristic Sizes of Life in the Oceans, from Bacteria to Whales. Annual Review of Marine Science 8:217–241.

[23] Pascual M, Dunne JA (2005) Ecological Networks: Linking Structure to Dynamics in Food Webs (Oxford Univ. Press).

[24] Trebilco R, Baum JK, Salomon AK, Dulvy NK (2013) Ecosystem ecology: size-based constraints on the pyramids of life. Trends in Ecology & Evolution 28:423–431.

[25] Fick SE, Hijmans RJ (2017) WorldClim 2: new 1-km spatial resolution climate surfaces for global land areas. International Journal of Climatology 37:4302–4315.

[26] Barneche DR, et al. (2021) Warming impairs trophic transfer efficiency in a long-term field experiment. Nature 592:76–79.

[27] McCauley DJ, et al. (2018) On the prevalence and dynamics of inverted trophic pyramids and otherwise top-heavy communities. Ecology Letters 21:439–454.

[28] McCann KS, Rasmussen JB, Umbanhowar J (2005) The dynamics of spatially coupled food webs. Ecology Letters 8:513–523.

[29] Mourier J, et al. (2016) Extreme Inverted Trophic Pyramid of Reef Sharks Supported by Spawning Groupers. Current Biology 26:2011–2016.

[30] Barneche DR, Allen AP (2018) The energetics of fish growth and how it constrains food-web trophic structure. Ecology Letters 21:836–844.

[31] White EP, Ernest SKM, Kerkhoff AJ, Enquist BJ (2007) Relationships between body size and abundance in ecology. Trends in Ecology & Evolution 22:323–330.

[32] Pearl J (2009) Causality: Models, Reasoning, and Inference (Cambridge Univ. Press).

[33] Shipley B (2016) Cause and Correlation in Biology (Cambridge University Press).

[34] Reicehnbach H (1956) The direction of time (The University of California Press).

[35] Bareinboim E, Pearl J (2016) Causal inference and the data-fusion problem. PNAS 113:7345–7352.

[36] Imbens GW (2020) Potential outcome and directed acyclic graph approaches to causality: Relevance for empirical practice in economics. J. of Economic Literature 58:1129–1179.

[37] Robinson JPW, Wilson SK, Jennings S, Graham NAJ (2019) Thermal stress induces persistently altered coral reef fish assemblages. Global Change Biology 25:2739–2750.

[38] Williams RJ, Martinez ND (2000) Simple rules yield complex food webs. Nature 404:180–183.

[39] Berlow EL, et al. (2009) Simple prediction of interaction strengths in complex food webs. Proceedings of the National Academy of Sciences 106:187–191.

[40] Barnes C, Maxwell D, Reuman DC, Jennings S (2010) Global patterns in predator–prey size relationships reveal size dependency of trophic transfer efficiency. Ecology 91:222–232.

[41] Reuman DC, et al. (2009) in Advances in Ecological Research (Academic Press) Vol. 41, pp 1–44.

[42] Nakazawa T, Ohba Sy, Ushio M (2013) Predator–prey body size relationships when predators can consume prey larger than themselves. Biology Letters 9:20121193.

[43] Edgar GJ, et al. (2020) Establishing the ecological basis for conservation of shallow marine life using Reef Life Survey. Biological Conservation 252:108855.

[44] Mora C, et al. (2011) Global Human Footprint on the Linkage between Biodiversity and Ecosystem Functioning in Reef Fishes. PLOS Biology 9:e1000606.

[45] Saha K, et al. (2018) The Coral Reef Temperature Anomaly Database (CoRTAD) Version 6 - Global, 4 km Sea Surface Temperature and Related Thermal Stress Metrics for 1982 to 2019. NOAA National Centers for Environmental Information. Dataset.

[46] Saavedra S, Bartomeus I, Godoy O, Rohr RP, Zu P (2022) Towards a system-level causative knowledge of pollinator communities. Philosophical Transactions of the Royal Society B: Biological Sciences 377:20210159.

[47] Edgar GJ, et al. (2014) Global conservation outcomes depend on marine protected areas with five key features. Nature 506:216–220.

[48] Karpowicz M, et al. (2021) Transfer efficiency of carbon, nutrients, and polyunsaturated fatty acids in planktonic food webs under different environmental conditions. Ecology and Evolution 11:8201–8214.

[49] Eddy TD, et al. (2021) Energy Flow Through Marine Ecosystems: Confronting Transfer Efficiency. Trends in Ecology & Evolution 36:76–86.

[50] Slobodkin LB (2001) The good, the bad and the reified. Evolutionary Ecology Research 3:91–105.

[51] Pozas-Schacre C, et al. (2021) Congruent trophic pathways underpin global coral reef food webs. Proceedings of the National Academy of Sciences 118.

[52] Small C, Nicholls RJ (2003) A Global Analysis of Human Settlement in Coastal Zones. Journal of Coastal Research 19:584–599.

[53] Halpern BS, et al. (2015) Spatial and temporal changes in cumulative human impacts on the world’s ocean. Nature Communications 6:7615.

[54] Hoegh-Guldberg O (1999) Climate change, coral bleaching and the future of the world’s coral reefs. Marine and Freshwater Research 50:839–866.

[55] Mora C, et al. (2006) Coral Reefs and the Global Network of Marine Protected Areas. Science 312:1750–1751.

[56] Moreno-Mateos D, et al. (2017) Anthropogenic ecosystem disturbance and the recovery debt. Nature Comm. 8:14163.

[57] Tabi A, Gilarranz LJ, Saavedra S (2022) Data from: Marine protected areas maintain pyramid-like structures of coral-reef fish communities, Dryad, Dataset.

[58] UNEP-WCMC, WorldFish Centre, WRI, TNC (2021) Global distribution of warm-water coral reefs, compiled from multiple sources including the Millennium Coral Reef Mapping Project. Version 4.1. Includes contributions from IMaRS-USF and IRD (2005), IMaRS-USF (2005) and Spalding et al. (2001). Cambridge (UK): UN Environment World Conservation Monitoring Centre.

[59] UNEP-WCMC and IUCN (2022) Protected Planet: The World Database on Protected Areas (WDPA) and World Database on Other Effective Area-based Conservation Measures (WD-OECM), January 2022, Cambridge, UK: UNEP-WCMC and IUCN.

[60] Center for International Earth Science Information Network CIESIN Columbia University (2020) Gridded population of the world, version 4 (gpwv4): Basic characteristics, revision 11. pal-isades,448 ny: Nasa socioeconomic data and applications center (sedac), (Gland, Switzerland: IUCN). Technical Report No. 26.

[61] Havens K (1992) Scale and Structure in Natural Food Webs. Science 257:1107–1109.

[62] Hamilton SL, et al. (2007) Size-selective harvesting alters life histories of a temperate sex-changing fish. Ecological Applications: A Publication of the Ecological Society of America 17:2268–2280.

[63] Kalisch M, Mächler M, Colombo D, Maathuis MH, Bühlmann P (2012) Causal Inference Using Graphical Models with the R Package pcalg. Journal of Statistical Software 47:1–26.

[64] Hopf JK, Caselle JE, White JW (2021) Recruitment variability and sampling design interact to influence the detectability of protected area effects. Ecological Applications.

[65] Froese, R. and D. Pauly. Editors. (2021) FishBase. World Wide Web electronic publication.

